# Re-membering: spontaneous reactivation of motor cortex during memory re-experiencing

**DOI:** 10.64898/2026.01.29.702278

**Authors:** Juliette Boscheron, Pepijn Schoenmakers, Arthur Trivier, Florian Lance, Bruno Herbelin, Dimitri Van De Ville, M. Babo-Rebelo, Olaf Blanke

## Abstract

Episodic autobiographical memory (EAM) relies on reactivations of cortical regions, originally engaged during event encoding. While reinstatement of auditory and visual regions during retrieval of external signals is well documented, reinstatement of motor regions, reflecting internal self-related signals from the person’s body, remains unexplored. Here, we investigated motor reinstatement at both cortical and peripheral levels. Using mixed-reality, participants encoded lifelike episodes involving specific motor actions. The next day, they freely retrieved these episodes while undergoing fMRI (N=30) or EMG (N=23). fMRI results revealed lateralized, effector-specific reactivation in primary and supplementary motor cortices. Moreover, hippocampal–motor connectivity increased during retrieval, and activity in premotor and supplementary motor areas scaled with subjective re-experiencing ratings. EMG recordings further showed, during retrieval, sub-threshold activation of the muscle active during encoding. Together, these results provide evidence for cortical and peripheral motor reinstatement during the re-experiencing of past events, emphasizing the embodied nature of EAM.

## Introduction

Episodic autobiographical memory (EAM) is characterized by the ability to travel back in time and re-experience episodes from one’s own life. EAM has been argued to be important for the feeling of being a unified and continuous self ^1,2^, allowing us to store and revisit personal past experiences by subjectively re-experiencing the associated contextual, emotional, and sensorimotor details of the event ^3^. Neuroimaging studies have consistently implicated the hippocampus as a central structure in EAM, due to its critical role in scene construction and mental time travel; i.e., the ability to disengage from the present and mentally reconstruct past or anticipate future events ^4–8^. However, while the hippocampus is essential for the reconstruction process of EAM, hippocampal activity alone does not explain the richness and modality-specific content of recollection ^9–12^.

During memory retrieval, the hippocampus interacts with neocortical regions that were originally active at the time of encoding, thereby enabling the reconstruction and re-experiencing of the event ^4,13,14^. This reconstruction process gives rise to reinstatement effects at both the hippocampal and neocortical levels ^15–21^. Thus, sensory-perceptual recollection of encoding-related information during retrieval has been associated with activation in visual ^10,15,17,22^ and auditory brain regions ^17,18,20^. The unfolding of these reinstatement effects has been argued to begin in ventro-medial prefrontal cortex (vmPFC) and medial temporal lobe (“access” phase), followed by a reactivation of neocortical sensory regions (“elaboration” phase) ^4,10,23,24^. Moreover, hippocampal–neocortical functional connectivity is enhanced during episodic retrieval ^25–27^ and correlates with visual vividness ^26^ as well as item and spatial precision ^25^.

Despite these important findings, several key questions remain unanswered. While sensory aspects of the *external world* (i.e., visual and auditory signals) have been widely studied in episodic memory research, the role of *internal signals* (i.e., motor or somatosensory) from the person’s body during encoding of particular events has received considerably less attention. This oversight is particularly striking given that such bodily signals are present in most instances of daily life, unlike visual inputs, which may be reduced or absent during encoding; e.g., when events are experienced with closed eyes. Furthermore, motor signals are self-generated, therefore potentially relevant to the sense of re-experiencing past episodes. Accordingly, bodily signals, particularly motor cues, may contribute to EAM, and reinstatement effects may also involve the motor and/or somatosensory cortices. Prior work has shown that motor regions are reactivated during the explicit recall of motor actions ^28^ and during motor imagery ^29^, but whether the primary motor cortex (M1) and other motor-related regions (premotor cortex: PMC, supplementary motor area: SMA) are engaged in reinstatement processes during EAM retrieval, in the absence of explicit motor imagery, has never been demonstrated.

One key challenge in the study of EAM is to have participants encode events in ecologically valid but experimentally controlled settings. Most past research has relied on retrospective reports of real-life events ^7,10,30–32^. Although these events are truly “autobiographical”, the original encoding conditions are unknown, unmeasurable, and vary in remoteness. Conversely, laboratory episodes afford tightly controlled events, but are often less realistic and less relevant to participants, often differing markedly from real-life events in their neural substrates ^33,34^. Consequently, there is a growing need for ecological laboratory-encoded events to capture richer, more realistic encoding and retrieval memory processes. We here developed a novel, highly immersive 3D mixed-reality setting in which participants encoded new episodes under experimentally controlled conditions. Importantly, they could see their own body while performing specific motor actions. Then, 24h after encoding, participants returned to the lab and retrieved these events as vividly as possible while undergoing fMRI (N=30; Study 1) or EMG recordings (N=23, different participants; Study 2).

We investigated (1) whether reinstatement during EAM retrieval involves the motor cortex and reflects the motor actions performed at encoding; (2) how such motor cortex reinstatement is associated with hippocampal activity; (3) how motor cortex activity relates to the subjective re-experiencing of the encoded event; (4) whether motor reinstatement extends to the peripheral-muscular level. We hypothesized that M1, as well as premotor and supplementary motor cortices, would be reactivated during retrieval and that their connectivity with the hippocampus would be enhanced. We also hypothesized that motor reactivations would correlate with subjective re-experiencing of memories and generate sub-threshold muscular reactivations.

## Results

During the encoding session (Fig. 1A, Supplementary Video 1: https://mediaspace.epfl.ch/media/LNCO_ReMembering_Boscheron_2026/0_vvd1mwj0/49346), participants were immersed in a 3D mixed-reality setting designed to maximize the ecological validity and immersiveness of encoding experiences ^35,36^. They drove a car along a country road (see Fig. 1A, Methods, Supplementary Fig. 1A), where 40 different obstacles (i.e., complex objects; see below) appeared successively. The virtual car stopped in front of each obstacle, and participants were instructed to perform a predefined sustained motor action (see below) that displaced the obstacle to the side of the road. The car would then start again, and the participant would drive it to the next obstacle (Fig. 1A). Participants were informed that we were studying spatial navigation to create conditions for implicit encoding, thereby simulating the naturalistic, incidental nature of EAM encoding.

**Figure 1:**
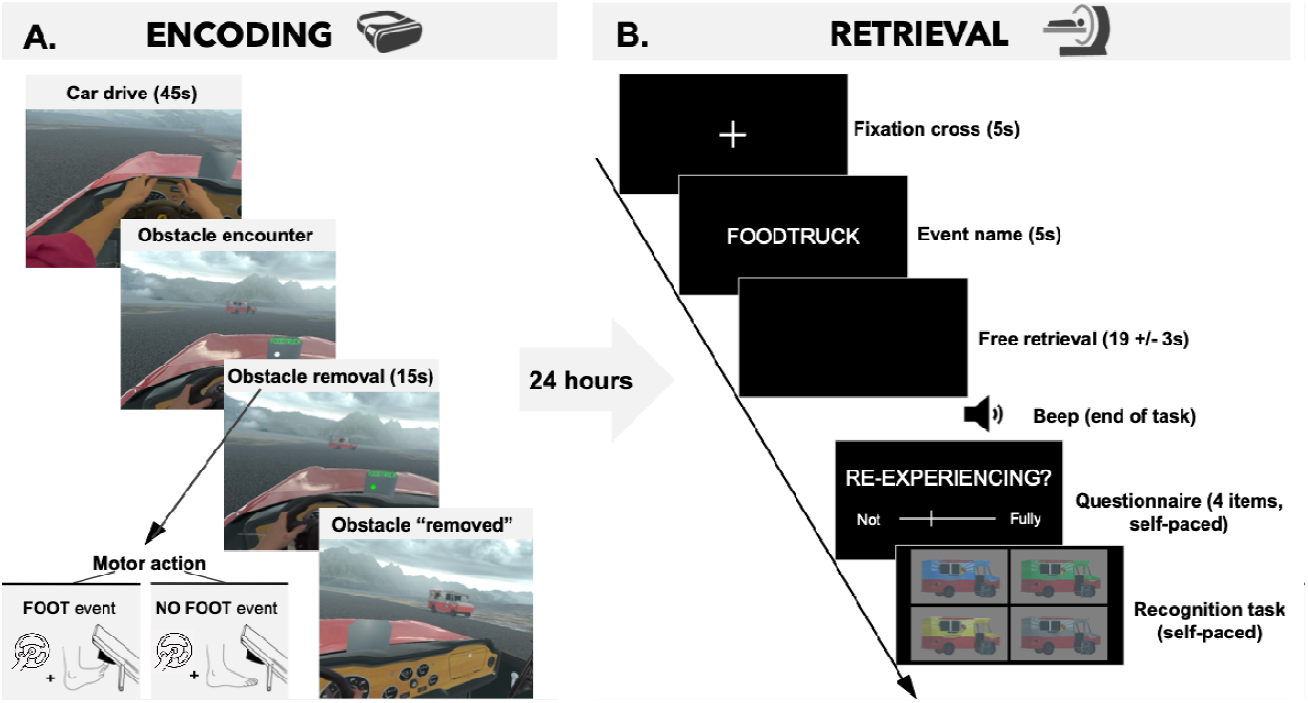
Experimental paradigm. **A**, Encoding session. Participants drove a car in a 3D mixed-reality environment and encountered obstacles that they had to move to the roadside, upon specific motor movements, depending on the condition: foot (button press and foot pedal lift) or no-foot (only button press). The animation of the obstacle moving away from the road constituted an event that participants implicitly encoded. The required motor action as well as the event name were indicated on the car dashboard. The pictures are screenshots taken from the performance of the task by one of the authors. **B**, Retrieval session, in the MRI scanner. In each trial, participants were shown the name of an event and were prompted to recall all the details of the scene and try to re-experience the event. After the “free retrieval” task, they had to rate their subjective experience on a set of Likert-scale questions (“Questionnaire”) and to recognize the obstacle amongst four color variations (“Recognition task”, 4-alternative forced choice).

Each obstacle was unique (e.g., a food truck; Supplementary Table 1) and was associated with a specific animation (e.g., a food truck packs away its signs, then pulls over to the right side of the road, Fig. 1A; a moose moves to the roadside, see Supplementary video 1), constituting an individual event. Following the dashboard instructions, participants performed one of two motor actions, constituting the two experimental conditions. The foot condition required a sustained right foot-pedal lifting engaging the right tibialis anterior muscle (foot events, N= 20), and the no-foot condition did not require any foot movement (no-foot events, N=20) (Fig. 1A, Supplementary Fig. 1B, and Methods for further details). Both conditions required a sustained left-thumb button press to maintain the same level of agency over the event unfolding for both conditions ^37^.

**Table 1:**
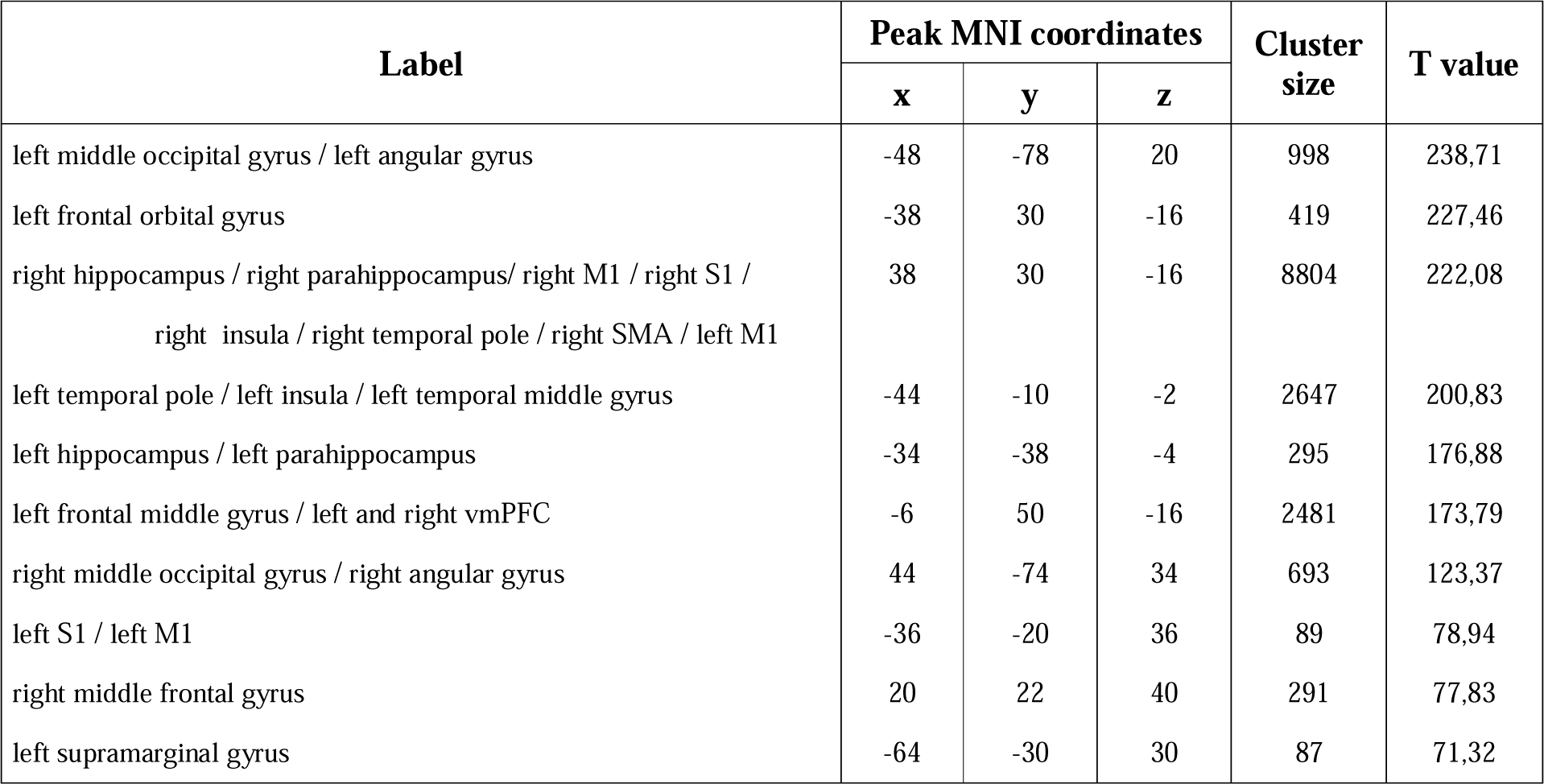
Main effect of Free Retrieval. Clusters showing higher BOLD activation during free retrieval task as compared to implicit baseline. Clusters are labeled using the Automated Anatomical Labeling (AAL) atlas ^41^. Reported coordinates correspond to MNI space.

| Label | Peak MNI coordinates |  |  | Cluster size | T value |
| --- | --- | --- | --- | --- | --- |
|  | x | y | z |  |  |
| left middle occipital gyrus / left angular gyrus | -48 | -78 | 20 | 998 | 238,71 |
| left frontal orbital gyrus | -38 | 30 | -16 | 419 | 227,46 |
| right hippocampus / right parahippocampus/ right M1 / right S1 /<br>right insula / right temporal pole / right SMA / left M1 | 38 | 30 | -16 | 8804 | 222,08 |
| left temporal pole / left insula / left temporal middle gyrus | -44 | -10 | -2 | 2647 | 200,83 |
| left hippocampus / left parahippocampus | -34 | -38 | -4 | 295 | 176,88 |
| left frontal middle gyrus / left and right vmPFC | -6 | 50 | -16 | 2481 | 173,79 |
| right middle occipital gyrus / right angular gyrus | 44 | -74 | 34 | 693 | 123,37 |
| left S1 / left M1 | -36 | -20 | 36 | 89 | 78,94 |
| right middle frontal gyrus | 20 | 22 | 40 | 291 | 77,83 |
| left supramarginal gyrus | -64 | -30 | 30 | 87 | 71,32 |

24 hours later, participants returned to the Lab to perform the retrieval session (Fig. 1B) in the MRI scanner (Study 1). Each trial focused on a single encoding event, prompted by the event name (e.g., “food truck”) presented on the screen for 5s, and involved three successive tasks. First, participants performed the free retrieval task, with the following instruction: “*Close your eyes and try to remember the obstacle moving in front of you and all the details of the scene, but also try to imagine yourself in the scene again, as if you were travelling back in time to re-experience or relive the scene*”. Second, participants evaluated different aspects of free retrieval: their sense of re-experiencing (“*To what extent were you re-experiencing this memory*?” with a response scale 1-7); their overall vividness for the memory, and the amount of visual details and bodily aspects they retrieved (Methods). Third, participants performed a recognition task, in which they were asked to identify the correct obstacle from encoding among four color variations.

On average, the 30 participants (16 females, 24.7 ± 3.4 years old) gave re-experiencing ratings of 4.31 (SD=0.85) on a scale ranging from 1 (not re-experiencing) to 7 (fully-re-experiencing), comparable to previous studies using real-life autobiographical events ^38–40^. More than half of the events were rated 5 or higher (Supplementary Fig. 2A). Participants performed above chance in the recognition task (mean accuracy=0.49, SD=0.12, chance level=0.25), indicating successful retention of the encoded content. Furthermore, correctly recognized obstacles were associated with higher ratings of re-experiencing (paired t-test t(29)=7.34, p<0.001, see Supplementary Fig. 2B). Finally, re-experiencing ratings did not differ between encoding conditions (paired t-test t(29)=−0.82, p=0.42, see Supplementary Fig.2C). These results validate the present task as an appropriate tool for the study of EAM retrieval under mixed-reality conditions.

**Figure 2:**
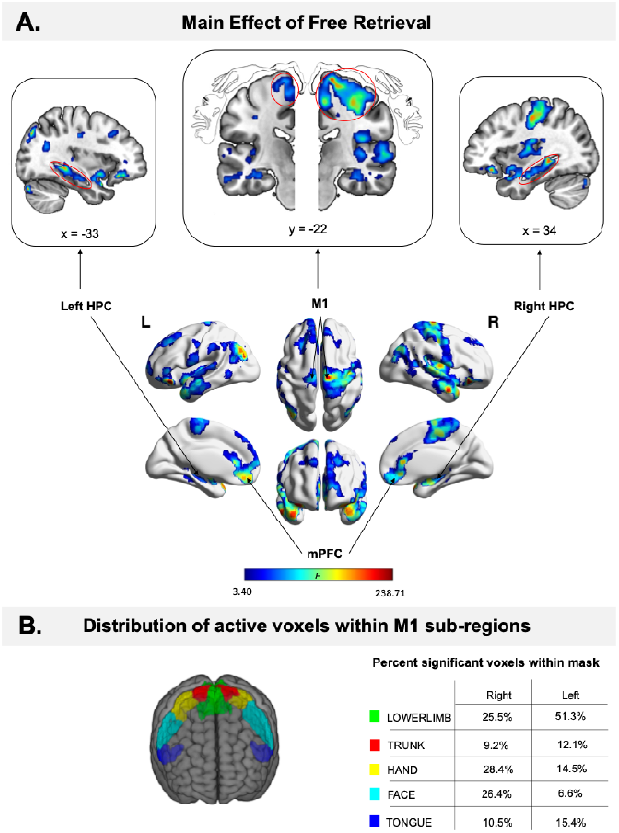
Motor cortex reinstatement during free retrieval. **A**, Main effect of free retrieval (p<0.001, FWE cluster correction). **B**, M1 subfield masks extracted from Brainnetome atlas and table of percentage of active voxels within each mask 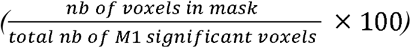 . M1: primary motor cortex, HPC: hippocampus, mPFC: medial prefrontal cortex, L: left, R: right.

### Motor cortex reinstatement during retrieval

fMRI analysis of the retrieval session (day 2) revealed several significant activation clusters during the period of free retrieval, during which participants, with eyes closed, mentally re-experienced the previously encoded events. Full brain analysis (free retrieval main effect, at the group level) revealed several significant activation clusters (Table 1), encompassing three classical EAM regions: bilateral hippocampus, bilateral parahippocampal gyrus, and vmPFC (Fig. 2A, see Table 1 for details of each cluster). We also observed activity in other regions that have been associated with episodic retrieval in previous studies; i.e., bilateral angular gyrus, bilateral middle temporal gyrus, bilateral insula, and bilateral middle and superior frontal gyrus (Fig. 2A).

Critically, motor regions including primary motor (M1) cortex and supplementary motor area (SMA), as well as primary somatosensory cortex (S1) were also activated (Fig. 2A). Moreover, the largest cluster in left M1, representing 51.3% of voxels in this region, was located within the lower limb area of left M1 (Fig. 2B), contralateral to the right foot that was engaged during encoding. Conversely, the largest cluster in right M1 was centered in the hand region (28.4% of right M1 voxels fell in the hand mask, Fig. 2B), contralateral to the left thumb engaged during encoding. Notably, during free retrieval, participants were never instructed to re-enact the motor actions they performed during encoding.

These results indicate that, in addition to classical memory structures (hippocampus, parahippocampal cortex, vmPFC), (1) motor cortex, consisting of M1 and SMA, is activated during free retrieval; (2) the observed M1 activation is in line with the specific motor actions that participants performed during encoding, in terms of laterality and somatotopy (left M1 foot region and right M1 hand region), consistent with motor cortex reinstatement during EAM retrieval.

### Somatotopic motor reinstatement in primary motor cortex

Next, we investigated whether motor reinstatement was specific to the motor actions performed during encoding. To this aim, we focused on brain activity in the foot area of left M1 (i.e., corresponding to the right foot pedal movement performed at encoding) during the retrieval of foot vs. no-foot events. Subject-specific regions of interest (ROIs) were defined using a “foot localizer” task, in which participants alternated between rest and right foot lifting; i.e., right tibialis anterior muscle contractions (Methods). Individual ROI masks were derived from a movement > rest contrast (Fig.3A). For each subject, we then extracted the mean contrast estimates within the right foot ROI mask in left M1, during the entire free retrieval, separately for the foot and no-foot events. This analysis revealed a trend for higher activity during the retrieval of foot vs. no-foot events (paired t-test: t(28)=1.74, p=0.093). To further explore this effect, we divided the 16-second free retrieval period in two 8s segments, considering a first 8s-access phase and a second 8s-elaboration phase, as done previously ^10,42,43^. Mean activity in the ROI was significantly different between retrieval of foot vs. no-foot events during the elaboration phase (Fig.3B, paired t-test: t(28)=2.51, p=0.036, Bonferroni corrected for the 2 phases tested), but not during the access phase (paired t-test: t(28)=-0.24, p=1, Bonferroni corrected). Critically, there was no significant difference in any of the analyzed time periods in the corresponding right M1 region, obtained by projecting the left M1 to the right hemisphere (paired t-test whole retrieval period: t(28)=0.75, p=0.46; paired t-test access phase: t(28) =-0.89, p=0.76, Bonferroni corrected for the two sub-phases tested; paired t-test elaboration phase: t(28)=1.98, p=0.116, Bonferroni corrected). This finding reveals a condition-specific somatotopic M1 activation.

After the retrieval session, we asked participants to indicate whether each event was associated or not with a foot movement during encoding. Participants’ responses were at chance level (paired t-test: t(29)=1.57, p=0.13) (Fig. 3C). Therefore, action-specific somatotopic M1 activation was implicit and not caused by intentional motor imagery of the foot movement.

**Figure 3:**
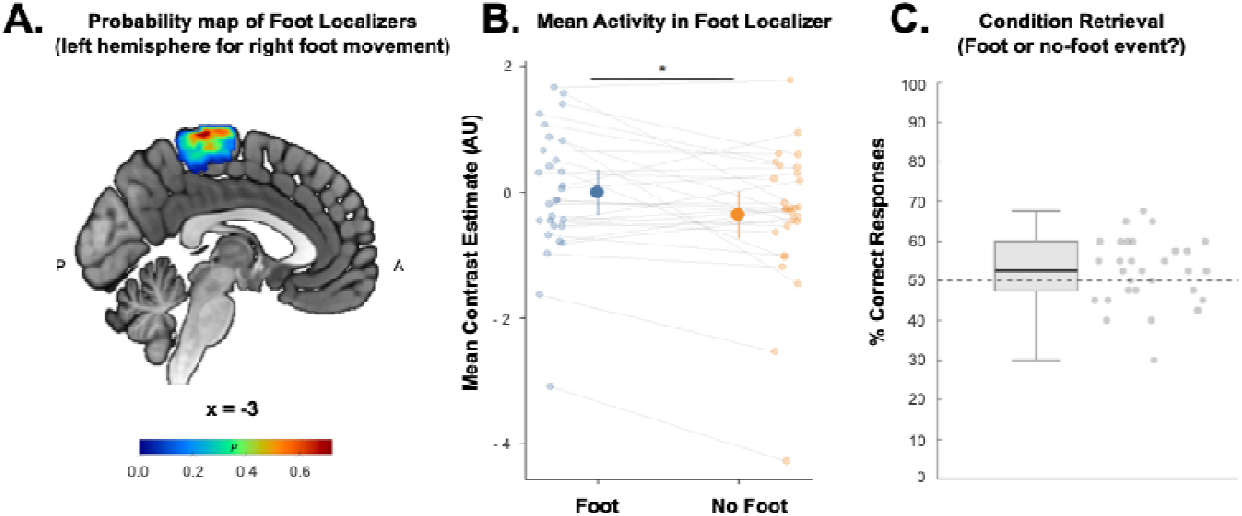
Activity in contralateral M1 foot region is higher during retrieval of foot events compared to no-foot events. **A**, Group-level probability map of subject-specific left M1 localizer masks (right foot movement). **B**, Mean contrast estimate in subject-specific left M1 localizer masks for retrieval of foot and no-foot events, during elaboration phase (8 to 16s of free retrieval). Each individual point depicts the mean contrast estimate value (arbitrary units) for a participant in a given condition, within their individual left M1 localizer mask. The two values of the same participant are connected with a straight line. **C**, Mean success rate to the question “Did this event involve a foot movement yesterday?”. The dashed line indicates chance level. *: p<0.05. A: anterior, P: posterior.

Taken together, these results reveal action-specific M1 reactivation during retrieval; i.e., increased activity in a specific body-part representation of M1 depending on whether that body part had been used during encoding. This was observed especially during the elaboration phase of event retrieval, absent in contralateral M1, and not associated with intentional motor imagery, which is compatible with motor reinstatement in EAM.

### Hippocampal functional connectivity with M1, premotor cortex and supplementary motor area increases during free retrieval

Next, we explored hippocampal functional connectivity (FC) across the entire brain during free retrieval to test whether motor regions would appear among the regions showing enhanced coupling with the hippocampus. We applied seed-to-whole-brain generalized psychophysiological interaction (gPPI) analyses ^44^, using the hippocampus of each hemisphere from our initial activation map as seed regions, and the main effect of free retrieval as a modulator. These analyses revealed significant task-related increase in FC between both hippocampi and different motor cortices, relative to the implicit baseline (Supplementary Table 2 for details of each cluster). For the right hippocampus, these included bilateral M1 and S1 as well as left SMA and right PMC (Fig. 4). The right hippocampus also showed task-related increase in FC with bilateral precuneus, bilateral posterior parietal cortex, bilateral primary visual cortex and left vmPFC. The same analysis for left hippocampus also showed enhanced FC with motor cortices in right M1, left PMC, right SMA and right S1 (Fig. 4). There was also a significant increase in FC between the left hippocampus and bilateral precuneus, bilateral posterior parietal cortex, bilateral primary visual cortex. When comparing hippocampal FC during retrieval of foot vs. no-foot events, we found no cluster that survived statistical corrections for multiple comparisons (all p > 0.05 after cluster-size p-FWE-correction).

**Figure 4:**
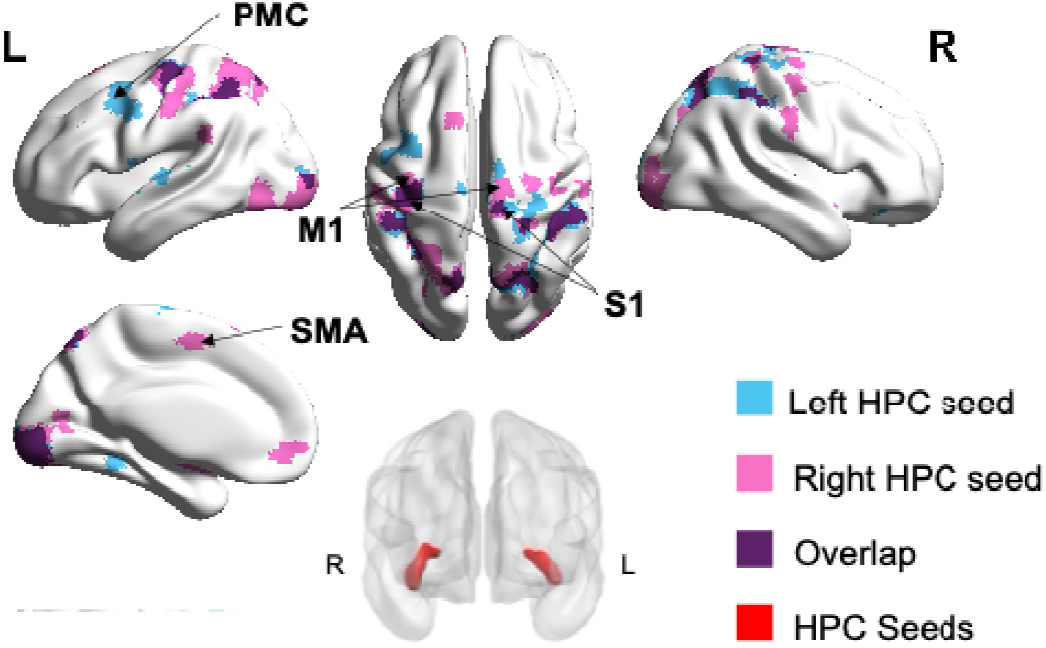
Hippocampus-to-whole-brain functional connectivity (FC) increases during free retrieval. Seed-to-whole-brain analysis of hippocampus FC during free retrieval (gPPI analysis). The right and left seeds considered for the analysis (red) were extracted from the main activation map of free retrieval (Fig.2). They correspond to activation clusters spanning from hippocampus to parahippocampus. Results from the gPPI analyses taking the right HPC and the left HPC as seeds are depicted in pink and blue, respectively. The overlap between significant clusters in the two analyses (left and right hippocampal seeds) is represented in purple. PMC: premotor cortex; SMA: supplementary motor area; M1: primary motor cortex; S1: primary somatosensory cortex; L: left; R: right.

These results show that both hippocampi enhance their functional connectivity with M1, PMC and SMA during retrieval (Fig.4), compatible with motor reinstatement in three key motor regions.

### Activity in premotor cortex and supplementary motor area correlates with re-experiencing intensity during retrieval

Next, we investigated whether brain activity during free retrieval was modulated by the subjective report of re-experiencing the encoded event. Whole-brain parametric modulation analysis (trial-by-trial re-experiencing ratings as a regressor) identified six clusters in which BOLD activity was positively modulated by the intensity of re-experiencing; i.e., the higher the re-experiencing, the higher the activity in these regions (Fig. 5). These clusters included bilateral temporo-parietal cortex (left supramarginal gyrus, right angular gyrus) as well as right inferior frontal gyrus, which have previously been shown to be involved in re-experiencing episodic events ^12,45^. Crucially, our analysis also revealed positive modulation in several motor regions. These were right PMC and bilateral SMA (Supplementary Table 3).

**Figure 5:**
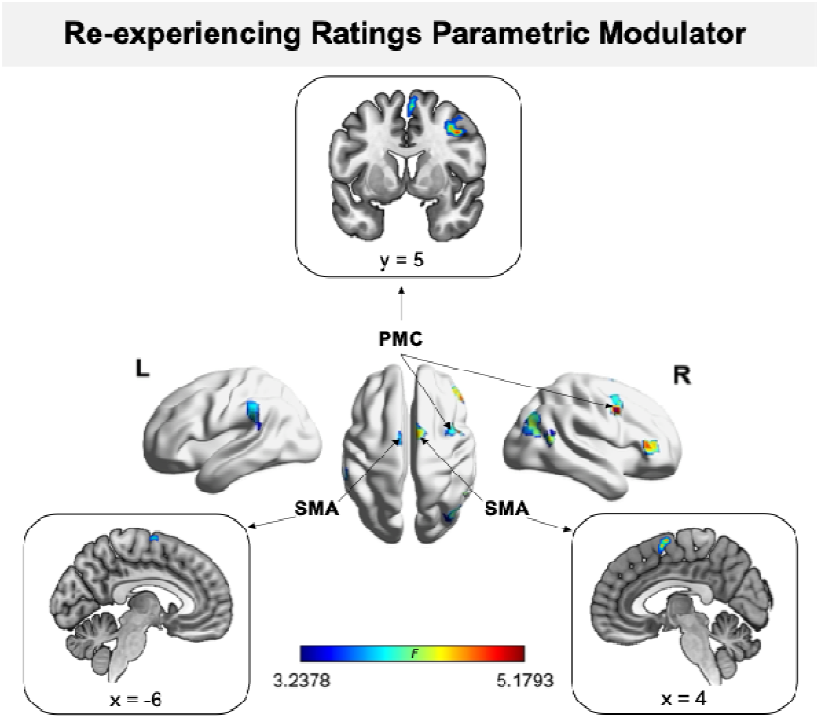
Premotor cortex and supplementary motor area activity positively correlate with re-experiencing intensity. Main effect of re-experiencing parametric modulator (p<0.001, FWE cluster correction). See Supplementary Table 3 for more details about the clusters and statistics. SMA: Supplementary Motor Area; PMC: Premotor Cortex; SMG: Supramarginal gyrus; L: Left; R: Right.

We observed no significant cluster in M1 or S1 (Fig. 5). These data suggest that higher-level motor cortex (PMC, SMA) is part of a re-experiencing network that also includes angular gyrus, supramarginal gyrus and inferior frontal gyrus.

### Electromyographic activity supports peripheral motor reinstatement (Study 2)

In Study 2, we investigated whether motor reinstatement in EAM could also manifest at the peripheral muscle level and be reflected in effector-specific changes in electromyographic (EMG) activity ^46–48^. For this, we tested a new group of 30 participants (Study 2), who performed an adapted version of the paradigm of Study 1 (Fig. 6A), where EMG from left and right tibialis anterior (TA) muscles was recorded during retrieval (Fig. 6B). Given that the right TA was actively engaged in pedal lifting during the encoding of foot events, and that the left TA was not engaged in any of the conditions, we predicted increased EMG activity for the right vs. left TA, during retrieval of foot as compared to no-foot events, in the absence of any overt movement. EMG data from seven participants were excluded (see Methods), leaving 23 participants in the final EMG analyses.

**Figure 6:**
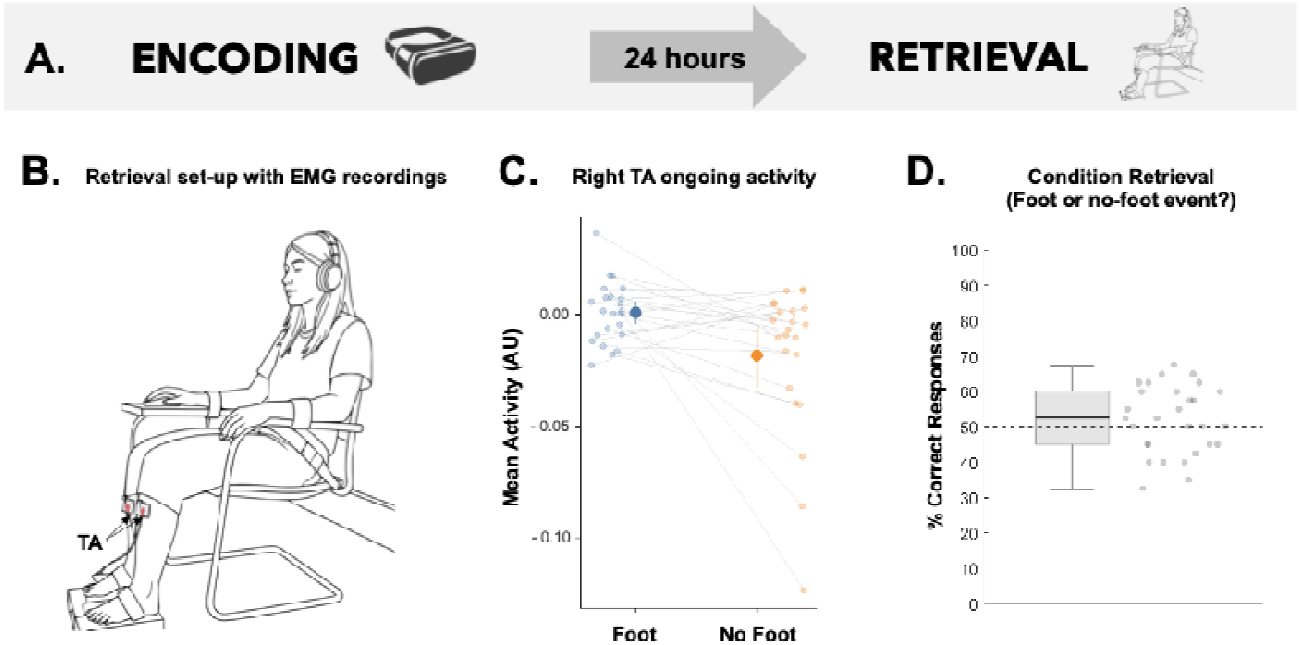
EMG motor reinstatement during retrieval (Study 2). **A**, Experimental design was similar to the fMRI study, with a mixed-reality encoding session on day 1 and a retrieval session 24 hours later. A new group of participants was tested. **B**, Bilateral Tibialis Anterior (TA) surface EMG activity was recorded while participants re-experienced previously encoded events. Participants were required not to move during the whole retrieval session. **C**, Mean EMG activity across the whole free retrieval in right TA for foot and no-foot events retrieval. Each individual point depicts the mean EMG activity (arbitrary units, AU) in right TA for a single participant. The two means of a same participant are connected with a straight line. **D**, Mean success rate to the condition retrieval question (“Did this event involve a foot movement yesterday?”). Success rate did not differ from chance (dashed line). **: p<0.005.

Retrieval of events involving a right foot movement at encoding (foot events) elicited higher EMG activity in the right TA than no-foot events (EMG averaged over the free reliving period; paired t-test: t(22)=3.04, p=0.006). This effect was significant during both the access (0-8s of free retrieval; paired t-test: t(22)=2.88, p=0.009 uncorrected) and the elaboration phase (Fig. 6C, 8-16s of free retrieval; paired t-test: t(22)=2.65, p=0.015 uncorrected). Critically, EMG did not differ between foot vs. no-foot events in the contralateral TA (left TA) in any of these time-windows (whole retrieval period: t(22)=-1.15, p=0.26; access phase: t(22)=-1.56, p=0.12; elaboration phase: t(22)=-0.31, p=0.76). EMG activity in the right tibialis anterior did not predict subjective re-experiencing ratings (main effect of right EMG activity: β=2.43, 95% CI [−1.63, 6.49], t(554.63)=1.18, p=0.24), regardless of encoding conditions (interaction right EMG activity * encoding condition: β=−0.70, 95% CI [−6.12, 4.72], t(553.11)=−0.25, p=0.80).

Importantly, EMG data were baseline-corrected relative to the fixation cross period, a period during which participants may exhibit some tension related to the expectancy of the upcoming retrieval task. Therefore, the EMG decrease observed in the no-foot condition for the right TA (Fig. 6B) may reflect progressive muscle relaxation as the trial unfolds. The right TA shows instead sustained EMG activity for the foot condition, supporting the idea that retrieval of foot events elicits sustained activity in those muscles involved in the encoded event, without any overt movement. Consistent with the MRI group, participants were at chance when asked whether an event was associated or not with a foot movement (mean accuracy=0.52, SD=0.12; one-sample t-test against chance level: t(22)=1.09, p=0.29, Fig.6D), excluding an effect of intentional motor imagery.

The results of Study 2 show, in a new sample of participants, that a muscle that had been active during encoding is reactivated during retrieval of the event. This effect was observed during both the elaboration and access phase of retrieval, was absent in the contralateral control muscle, and was not related to intentional motor imagery. Accordingly, these data show motor reinstatement effects in the peripheral motor system.

## Discussion

With a novel mixed-reality paradigm, we created an ecologically valid, highly immersive, and experimentally controlled setting for participants to encode new memories, involving specific motor actions. To probe the existence of motor reinstatement during EAM retrieval, we captured motor reactivations in the motor cortex (fMRI, Study 1) and in the peripheral motor system (EMG, Study 2). Alongside the network of classical memory regions, participants reactivated motor regions such as M1 and SMA, when retrieving previously encoded events. M1 activation was effector-specific, showing a lateralized somatotopic distribution, consistent with the motor actions performed at encoding. In addition, the hippocampus showed increased functional connectivity with M1, PMC, and SMA. Taken together, these results are compatible with *cortical motor reinstatement* in three key motor regions. Furthermore, we saw that activity in PMC and SMA was modulated by re-experiencing ratings, suggesting that higher motor reinstatement is associated with better re-experiencing. In a separate EMG study with a new sample of participants, we extended these results. During retrieval, we observed enhanced activity in the muscle engaged during encoding, consistent with *peripheral motor reinstatement*. None of these effects can be explained by intentional motor imagery, as participants of both studies were unable to explicitly associate the events with the motor actions performed at encoding. Collectively, these results provide robust evidence for cortical and peripheral motor reinstatement in EAM, associated with the subjective sense of reliving past events.

Using fMRI, we observed robust engagement of classical EAM regions during re-experiencing, including the medial temporal lobe and medial prefrontal cortex, consistent with previous imaging findings ^21,24,49^. Activity was also enhanced in the angular gyrus, supramarginal gyrus, and precuneus, regions that have been repeatedly shown to be involved in EAM ^12,50–52^. Critically, we also observed significant activation in the primary motor cortex (M1) during memory retrieval, in line with our initial hypothesis. We also reported activation in primary somatosensory cortex (S1) and supplementary motor area (SMA) during retrieval, consistent with the fact that most movements involve not only M1, but also other motor and somatosensory regions. Indeed, movements are always accompanied by changes in proprioceptive and tactile input ^53^ and involve mechanisms of motor selection, planning, and coordination ^54^. Rather than being a general and unspecific bodily reinstatement, the present M1 activation cluster also shows somatotopic consistency ^55^, which is compatible with the specific body parts involved in the motor actions participants performed during encoding; i.e., the left hand and/or the right foot. This specificity can be paralleled with results from visual reinstatement showing different patterns of activity during retrieval of different categories of visual stimuli, such as faces or locations ^56^.

This lateralized limb-specific motor reinstatement effect was most prominent during the second half of the free retrieval period, consistent with previous findings showing that both subjective re-experiencing and neural reinstatement of encoding-related activity emerge predominantly during the elaboration phase of EAM retrieval ^10,42^. These effects occurred while participants were unable to explicitly report which events had been associated with a foot action at encoding. Of note, the thumb and foot actions performed during encoding were not attentionally demanding, and participants were more likely to attend to obstacles moving away than to these sustained, repetitive motor actions. These results are therefore markedly different from known motor reactivations occurring in M1, PMC and SMA during the explicit recall of actions ^28,57^ or during motor imagery ^58,59^. In the present study, the motor action is subsidiary to the encoded event and is automatically and implicitly reactivated during recall. This is in line with previous findings showing that both visual and auditory cortical reinstatement can occur under implicit memory conditions, in which participants are not instructed to engage in explicit visual or auditory imagery ^60–62^.

Previous research has failed to demonstrate M1 involvement during memory retrieval ^11,43,63^, which may be explained by several methodological factors. On the one hand, motor actions at encoding have never been a factor of interest, measured and/or accounted for with the appropriate control conditions, so as to reveal effector-specific retrieval effects. On the other hand, motor activity at retrieval has been treated as a confounding factor and discarded. In studies focusing on the retrieval of real-life autobiographical events, participants are often asked to press a button to signal that they have accessed a memory ^10,42^, or sometimes to orally describe the content of their retrieval process ^16^. In contrast, we chose to examine the main effect of free retrieval without a contrasting baseline, thereby avoiding such confounds.

Overall, the present results extend the concept of sensory reinstatement to the motor domain, as previously observed in the visual ^10,15,17,22^ and auditory ^17,18,20^ domains. We argue that cortical motor reinstatement shows properties that are similar to those observed for visual and auditory reinstatement: (1) specificity to the bodily effector (comparable to specificity to visual/auditory stimulus category), (2) predominance during the elaboration phase of retrieval, and (3) implicit nature of reinstatement. While sensory reinstatement reconstructs the sensory aspects of the scene, we here show that memory retrieval is also associated with the *re-embodiment of the event*, thereby reinstating the observer or subject in the encoded event.

In the present study, we also replicate previous findings showing enhanced hippocampal functional connectivity (FC) with prefrontal and posterior parietal areas during the retrieval process ^25–27^. Both hippocampi showed task-related enhanced FC with bilateral precuneus, bilateral posterior parietal cortex, and bilateral primary visual cortex, regions that have been shown to be involved in EAM retrieval ^10,45,64^ as well as bodily processing ^65,66^. Critically, we report a new finding of increased FC between right and left hippocampi and M1, S1, as well as PMC and SMA. This strengthened FC is also compatible with the mechanism of motor reinstatement and suggests an interaction between memory retrieval processes and a motor network composed of regions involved in movement execution, motor preparation, and coordination ^54,67^. Furthermore, this result is compatible with current views on the temporal unfolding of retrieval mechanisms, according to which the hippocampus drives the reinstatement of neocortical regions ^49^. Therefore, motor reinstatement might be triggered by the hippocampus, via increased hippocampal-motor network connectivity, but further investigation is needed to address this point specifically.

In Study 2, we tested a new group of participants and showed that motor reinstatement goes beyond the motor cortex, eliciting limb-specific increases in muscular activity. Specifically, we found increased EMG activity in the right tibialis anterior muscle during the retrieval of events that had required that muscle for the right foot movement at encoding. This muscular “reactivation” was below threshold (i.e., no overt movement was produced) and was implicit (i.e., it emerged despite participants’ inability to explicitly report which events had been associated with a foot movement during encoding). Previous work showed that EMG activity is modulated during explicit motor imagery (i.e., ^68^), in the absence of overt movement. The present data suggest that EMG activity is a marker of memory retrieval. Critically, the left tibialis anterior control muscle (i.e., not involved in movement during encoding) did not show any EMG modulation at retrieval. The laterality and limb-specificity are consistent with the fMRI findings showing the somatotopic and lateralized organization of the motor system, and supporting the idea that memory retrieval re-engages or “re-members” the same neural regions and limb muscles involved at encoding.

While cortical motor reinstatement occurred predominantly during the elaboration phase of memory retrieval, the EMG reactivation occurred during both the access and elaboration phases. One possible explanation could be the body posture congruency between encoding and retrieval sessions in the EMG study, as participants sat in a similar posture in both sessions, whereas in Study 1, participants sat during encoding and lay down in the MRI scanner during retrieval. Congruent body postures have been shown to facilitate memory retrieval ^69–71^, and this factor may facilitate motor system reinstatement. Future work, directly testing this fascinating potential postural effect in motor reinstatement, is needed.

The present data show that motor reinstatement is also of relevance for conscious elements of event retrieval. Increased visual reinstatement has previously been associated with higher vividness ratings ^10,15,72^. Here, we extend this finding to motor reinstatement and the phenomenology of event retrieval, showing that increased motor reinstatement was associated with greater event re-experiencing. Specifically, activity in the SMA and PMC, but not in M1, was positively modulated by the participants’ ratings of re-experiencing. These higher-order motor areas are essential for movement generation and planning, with the SMA primarily involved in internally generated movement planning, coordination of complex or sequential actions, and bilateral motor control ^54^ and the PMC in externally guided movement selection, sensory integration, and preparation for voluntary actions ^54,67^. Based on our results, SMA and PMC consistently contribute to the motor dimension of EAM retrieval, as they appeared in all our fMRI analyses; i.e., including motor reinstatement mechanisms, hippocampal connectivity, and the association with subjective experience during memory retrieval. On the contrary, while M1 consistently showed reinstatement effects, its activity did not relate to re-experiencing. This suggests that more global bodily representations in SMA and PMC, but not the limb-specific M1 activations, contribute to subjective aspects of EAM, further supported by our finding that peripheral limb-specific motor reinstatement was also not correlated with re-experiencing of the events.

Beyond these motor regions, we observed a positive correlation between re-experiencing ratings and activity in the right angular gyrus, left supramarginal gyrus, and precuneus, a set of regions consistently implicated in bodily self-consciousness and retrieval vividness ^10,45,50,64,65,73^. The right angular gyrus, was shown to not only support spatial bodily perception ^65^, but also to be involved in the subjective vividness and richness of autobiographical retrieval ^64,74^. The left supramarginal gyrus has been associated with proprioceptive integration and motor planning ^66^, as well as bodily self-consciousness ^75,76^, while the precuneus has been implicated in motor imagery and self-referential thought ^73^ as well as vividness of EAM retrieval ^10,64^. These converging findings support the view that memory re-experiencing is closely tied to the reactivation of body-centered and action-related neural processes, highlighting the embodied/re-membered nature of EAM retrieval. Each event that we encode in memory is perceived from within a body, and the constant integration of bodily-related sensory and motor signals at the time of encoding has been suggested to be important for the feeling of experiencing that event ^65,77^. Therefore, we argue that when participants later retrieve the event, the extent to which these bodily signals from encoding are reinstated contributes to shaping the subjective feeling of re-experiencing the memory of the event ^78^.

Several limitations of the present work should be acknowledged. First, although our mixed-reality paradigm increased ecological validity, the encoded events remained experimentally constrained and were neither part of our participants’ personal history nor autobiographically relevant. Motor reinstatement during retrieval of real autobiographical events may manifest differently, potentially also reflecting affective states and social interactions. Second, the temporal dynamics of motor reinstatement cannot be fully resolved with fMRI due to its limited temporal resolution. Third, while hippocampal–motor functional connectivity increased during retrieval, we cannot claim that motor reinstatement is initiated by the hippocampus. Future work combining connectivity modelling with temporally resolved methods (M/EEG) will be essential to further characterize the temporal and network dynamics of motor reinstatement.

To conclude, our findings provide robust evidence for motor reinstatement during memory retrieval at both central and peripheral levels. Primary (M1, S1) and higher-order (PMC, SMA) motor regions are differently involved in the reactivation process, with only the latter regions relating to the subjective feeling of re-experiencing the event. These results highlight the role of the brain representation of the body of the subject who encodes the events, by showing that multiple neural traces of the bodily movements performed at encoding are stored and reactivated during retrieval. Motor systems are therefore inherently involved in the re-experiencing of past events, highlighting the embodied or re-membered nature of EAM. By showing that remembering is re-membering, the present work opens new avenues for understanding memory as a sensorimotor simulation process that dynamically engages the body to reconstruct and reenact our personal past.

## Methods

### Study 1

#### Participants

Thirty participants (right-handed, 16 females, 24.7 ± 3.4 years old) were recruited. They had normal or corrected-to-normal vision and no history of psychological nor neurological disorders. All participants provided informed written consent in accordance with the Cantonal Ethical Committee of Geneva (2015-00092) and the declaration of Helsinki (2013).

#### Experimental design

The experiment included two sessions separated by 24 hours, i.e. an encoding session (day 1) and a retrieval session (day 2) in the MRI scanner (Fig.1A).

Encoding Session (Day 1)

##### Set-up

The experiment took place in a dedicated room with walls of a uniform color, furnished with a table and a chair (Supplementary Fig.1). The table was equipped with the steering wheel and foot pedal of a car simulator (Thrustmaster T80 Ferrari 488 GTB Edition, Guillemot Corporation S.A, France; Supplementary Fig.1A). The foot pedal was mounted upside down to be activated by lifting-movement instead of pressing-movement of the foot. Participants’ right foot was placed exactly under the pedal so that they could activate it only with dorsiflexion, without displacing the leg. Participants were seated in front of the steering wheel at a comfortable distance that allowed for a natural reach (Supplementary Fig.1A). Participants were equipped with a Varjo XR3 (Varjo Technologies, Helsinki, Finland) mixed-reality headset with video see-through, allowing them to see their real body and the steering wheel integrated into the virtual environment. This headset has a human-eye resolution with 2 screens per eye; a central focus area (27° x 27°) at 70 PPD uOLED, 1920 x 1920 pixels, and a peripheral area at over 30 PPD LCD, 2880 x 2720 pixels, leading to a 115° horizontal field of view at a maximum refresh rate of 90Hz. It is equipped with 12-megapixel video pass-through cameras at 90 Hz, allowing for the mixed-reality integration of steering wheel and participant’s body inside the virtual car cockpit using chroma-keying (cutting out video content of the room based on color; Supplementary Fig.1B). White noise was diffused in participant’s ears throughout the whole encoding session via AKG headphones. The 3D virtual environment was implemented in Unity 3D (version 2019.4.31f, Unity Technologies, USA) with Unity Engine standard pipeline and Varjo plugin (version 2.2.0 for Unity XR SDK version 4.0.1). 3D models were designed in-house and customized to meet the specific requirements of the task.

##### Encoding paradigm

The virtual environment consisted of a country road surrounded by mountains (Supplementary Fig. 1B). Participants drove a car along the road. Participants were asked to maintain a “classic” driving posture throughout the task, with parallel feet on the ground and the two hands on the steering wheel in an academic driving position (Supplementary Fig.1A). They were free to move the car from right to left along the road using the steering wheel but had no control over the car’s speed, which was constant. After 45s of driving, an obstacle appeared on the road, blocking their way. The car stopped, the name of the obstacle (e.g., “Foodtruck”) appeared on the car’s dashboard, and participants had to perform specific motor actions to move the obstacle away (Fig. 1B). This constituted an event that participants implicitly encoded in memory. Each obstacle was associated with one of two possible action conditions, indicated by a symbol appearing on the dashboard. In the no-foot condition, a sustained left thumb button press was required, whereas in the foot condition a sustained left thumb button-press and a right foot-pedal lift were required. The animation was paused if participants interrupted the required motor action(s), and it continued only when they resumed performing the action(s). This obstacle removal phase lasted until the obstacle was completely out of the way (15s). The car would then automatically resume its course until the next obstacle (Fig.1B).

##### Experimental procedure

Participants were not informed that this was a memory task to avoid explicit memory encoding. They were told this study investigated the neural mechanisms of navigation and that the second day would consist of a debriefing about their VR experience. We provided standardized instructions to all participants, after which participants familiarized themselves with driving the car in the virtual environment and performing the motor actions. The training session included two events of obstacle removal, one in each of the two action conditions. The main task consisted of a sequence of 40 navigation-obstacle removal events, each event associated with one of the two conditions (20 foot events and 20 no-foot events) and a unique obstacle (see Supplementary Table 1 for full obstacle list). The occurrence of foot and no-foot conditions was pseudo-randomized, with a minimum of 2 and a maximum of 4 consecutive events of the same condition. After 20 events, participants took 2 to 5 minutes break during which they could remove the headset to rest their eyes and neck.

Retrieval Session (Day 2)

##### Set-up

Participants were laying in a 3T MRI scanner at Campus Biotech, Geneva. The retrieval task was developed with the in-house software ExVR (https://github.com/BlankeLab/ExVR) and displayed on a computer screen, which participants could see through a mirror reflection. Participants provided their responses using an MRI-compatible button box (Fiber Optic Response Pad, Current Designs) with their right hand.

##### Retrieval paradigm

The encoding session took place 24h after encoding. Each trial of the task was designed as follows (Fig.1C). First, a central fixation cross appeared for 5s. Then, a written cue was displayed for 5s, indicating the name of an obstacle from Day 1 to retrieve. Participants were asked to close their eyes and freely re-experience the event associated with this obstacle (see detailed instructions in Supplementary Box 1) for 19.5s±3s (randomized). An auditory cue then indicated the end of the free retrieval period, and participants were instructed to open their eyes. They were asked 4 self-paced questions about their subjective experience of retrieval of the event, derived from established autonoetic consciousness questionnaires (MCQ ^79^, EAMI ^31^). The first question probed the feeling of re-experiencing the event: “To what extent are you re-experiencing this memory?” (response scale: 1-No re-experiencing, 7-Fully re-experiencing) (EAMI ^31^). The second question examined the overall vividness of their memory of the event: “My memory for this event is” (response scale: 1-Dim, 7-Sharp/Clear) (MCQ ^79^). The third question examined the visual vividness of their memory of the event: “My memory for this event involves visual details” (response scale: 1-Little/None, 7-A lot) (MCQ ^79^). The last question probed the retrieval of bodily aspects of the event: “My memory for this event involves body movements” (response scale: 1-Little/None, 7-A lot). Finally, participants performed a recognition task where 4 color variations of the obstacle were presented. Participants selected the obstacle they believed they had seen at encoding (4-alternative forced choice). In study 2, participants also rated their confidence in their response to the recognition task (1-low, 7-high confidence).

##### Experimental procedure

Participants first completed a brief training session of the retrieval task using the training events from encoding, which allowed for any necessary clarification of the task instructions. Then, in the MRI scanner, participants performed two blocks of the retrieval task, each with 20 trials corresponding to 20 events, in random order. Finally, participants were shown all 40 obstacle names and reported whether each obstacle was associated with a foot movement or not (2-alternative forced choice).

#### MRI Data Acquisition

Structural and functional MRI data were acquired on a Siemens MAGNETOM Prisma 3T scanner equipped with a 64-channel head-and-neck coil at Campus Biotech Geneva (https://www.campusbiotech.ch/). T1-weighted anatomical images were acquired using a 3D MPRAGE sequence with the following parameters: repetition time (TR)=2200ms, echo time (TE)=2.98ms, inversion time (TI)=900ms, flip angle=9°, 208 sagittal slices, voxel size=1 × 1 × 1 mm3, and field of view (FOV)=256 × 256 mm2. GRAPPA acceleration (factor 2) was applied. Functional images were acquired using a T2*-weighted gradient-echo echo-planar imaging (EPI) sequence with multiband acceleration (factor 5). Acquisition parameters were: TR=1300ms, TE=32ms, flip angle=64°, 65 interleaved axial slices, voxel size=2 × 2 × 2 mm3, and FOV=224 × 224 mm2. The phase-encoding direction was posterior-to-anterior (j−), and partial Fourier (7/8) was used in the phase-encoding direction. Four dummy volumes were acquired prior to each run to allow for steady-state magnetization. A B□ field map was also acquired for distortion correction.

#### fMRI analyses

##### fMRI Preprocessing

Preprocessing was performed with SPM12 v7487 (http://www.fil.ion.ucl.ac.uk/spm) implemented in MATLAB R2021a (MathWorks). T1 images were defaced to remove facial features. Origin of functional images was reset to ensure consistency in orientation across subjects. Voxel displacement maps (VDM) were calculated using the field maps derived from the phase difference between two magnitude images acquired at different echo times (Short TE=4.92ms, Long TE=7.38ms). Anatomical scans were segmented into gray matter, white matter, and cerebrospinal fluid. Functional images were spatially realigned to the mean volume and unwarped using the previously calculated VDM, to correct for motion artifacts and distortions, respectively. Functional volumes were coregistered to anatomical volumes and further normalized to the MNI (Montreal Neurological Institute) standard space using the deformation fields obtained from anatomical segmentation. Finally, functional volumes were spatially smoothed using a Gaussian kernel of 4mm.

##### Univariate Analysis

The 1st and 2nd level General Linear Models (GLM) were estimated using SPM12 v7487 (http://www.fil.ion.ucl.ac.uk/spm) implemented in MATLAB R2021a (MathWorks). A 1st level GLM was implemented for each participant, with 12 regressors for each run (24 regressors in total): 2 “fixation cross” regressors, 2 “free retrieval” and 2 “re-experiencing parametric modulators” (one for foot and no-foot events each time) as well as 6 motion parameter regressors. For each participant, we computed two contrast images: one for free retrieval of foot events and one for no-foot events. At the second level, these contrast images were entered into a within-subject ANOVA, treating condition as a repeated-measures factor and subject as a random effect. The main effect of free retrieval was assessed by comparing the average foot and no-foot retrieval conditions against an implicit baseline across subjects (F-test). Next, to identify regions whose activity scaled with subjective re-experiencing ratings, we computed, at the first level, a parametric modulation contrast that collapsed foot and no-foot trials. These participant-specific parametric modulation contrast images were then entered into a one-sample t-test at the second level to assess significant modulation across participants. Effects were explored with a cluster-defining threshold of p<0.001 uncorrected, and an additional cluster-extent threshold determined using SPM’s random field theory (RFT)–based cluster-level family-wise error (FWE) correction at p<0.05, whole brain. The minimum cluster sizes required for significance were k=87 voxels for the main effect of free retrieval, and k=89 for the re-experiencing parametric modulator.

##### Foot and thumb localizers

At the end of the MRI task, participants performed two localizer runs, designed to identify the brain regions responsible for “right foot” and “left thumb” movements at the single-subject level. In each run, participants alternated between 20s rest and 20s of movement, following the instructions of a visual cue. During movement phases, participants were required to contract the target limb for about 2s, and gently release for about 2s. The right foot localizer engaged the right tibialis anterior muscle, while the left thumb localizer engaged the left adductor pollicis. To determine the individual M1 masks corresponding to right-foot and left-thumb movements, we contrasted “move” vs “rest” at the 1st level for each subject. We then extracted the 200 most active voxels (i.e., highest t-values) within a predefined global M1 mask, constructed as the sum of all M1 sub-regions from the Brainnetome atlas ^80^. Left foot masks were obtained by projecting the individual right foot masks onto the contralateral hemisphere and excluding overlapping voxels.

From the 1st level contrasts of the Free Retrieval GLM representing all foot and no-foot episodes respectively, we computed the mean contrast values across all voxels in each mask, per subject, leading to mean contrast values within the foot and thumb localizers, per subject, per condition.

##### Seed-to-whole-brain functional connectivity analysis

Connectivity was assessed using CONN toolbox 2021a (http://www.nitrc.org/projects/conn, RRID:SCR_009550) running in MATLAB R2018a (MathWorks). Preprocessed data were band-pass filtered from 0.008 to 0.9□Hz, and white matter, cerebrospinal fluid, and realignment parameters were regressed out. Subsequently, seed-to-whole-brain generalized psychophysiological interactions (gPPI) connectivity was extracted, with hippocampal masks as seeds of interest. The right and left hippocampus masks were derived from the univariate analysis of the main effect of free retrieval. Seed-to-whole-brain gPPI analysis allows us to study how the functional connectivity of a seed region with the rest of the brain is modulated by task events. Brain maps were thresholded using Random Field Theory parametric statistics as implemented in CONN toolbox with a threshold of p <0.05 cluster-size p-FWE-corrections.

#### Behavioral analysis

Statistical analyses were conducted using R (version 4.3.3). To compare subjective re-experiencing intensity between correctly and incorrectly recognized items, and between foot and no-foot events, we used paired t-tests across participants. To assess whether performance in identifying the encoding condition differed from chance level, we performed a one-sample t-test against chance (0.5). Normality assumptions were verified using Shapiro–Wilk tests. All tests were two-tailed, and significance was set at α=0.05.

### Study 2

#### Participants

The study included 30 participants (21 females, right-handed, mean age=25.37 years, SD=4.04 years), having normal or corrected-to-normal vision and no history of psychological disorders. Participants had not participated in the fMRI study and were naïve to its purpose. They were informed that the day 2 session was for debriefing. All participants provided informed written consent in accordance with the Cantonal Ethical Committee of Geneva (2015-00092) and the Declaration of Helsinki (2013).

Participants performed the same task as in Study 1. Two participants were excluded due to technical complications during recording. Another participant was excluded as he systematically answered 3 or lower on all behavioural questions, indicating he did not perform the retrieval task as expected. Four more participants were excluded due to an excessive number of artefacts on their EMG signal (>50% trials were discarded). The final sample included in the analyses was of N=23 participants.

#### EMG recordings and retrieval session set up

During the retrieval session, participants were equipped with bipolar EMG electrodes to record activity of the left and right tibialis anterior muscles (Noraxon Desktop DTS receiver coupled with their DTS Lossless EMG Sensors and proprietary myoRESEARCH software). The recording electrodes were positioned on the most proximal third of the tibialis anterior muscle, whilst the ground electrode was placed on the kneecap.

The retrieval session took place in a recording room protected from external electromagnetic noise. The task was displayed on a computer screen. Participants provided their responses on a keypad attached to the right armrest of the chair. The participants’ position was secured using several straps: a hip strap that pressed them into the seat, a strap around each forearm that fastened their arms to the armrests, and a strap around each foot that attached them to custom-made plastic receptacles.

#### EMG analyses

##### EMG preprocessing

EMG preprocessing and analysis were performed using the FieldTrip (fieldtrip-20240111) toolbox ^81^ on MATLAB (23.2.0.2428915). A bandstop filter was applied to remove power line noise (50Hz) and its harmonics. The data was segmented into epochs starting from -10s to 16s relative to the start of the free retrieval period.

Noise detection was performed visually for each individual trial, independently for each leg, using the Fieldtrip browser tool (ft_databrowser). The inspector was blind to the experimental conditions. We marked all segments of data containing noise; i.e., real movement, twitches, or any type of electrical artefact. The goal was to have only the ongoing, subthreshold activity of the tibialis anterior muscles. If the segments containing noise spanned either half or more of the fixation cross window ([-10 -5s]), or half or more of the entire trial, the trial was excluded.

If more than half of the trials were discarded for a given participant, the participant was excluded from analysis (n participants excluded = 4).

We considered the signal after full-wave rectification (i.e., absolute value). A trial-by-trial baseline correction was applied by subtracting the average activity obtained during the fixation cross window ([-10 -5s]) from the activity of the whole trial. Then, for each subject, EMG data were averaged over trials for each condition.

##### Statistical analysis

Paired t-tests (ttest function, Matlab) were performed to assess statistically significant differences in mean EMG activity between conditions across subjects. We verified normality for the EMG data (Shapiro-Wilk test, p=0.70, W=0.97).

#### Behavioral analyses

Statistical analyses were conducted using R (version 4.3.3). To examine whether trial-by-trial EMG activity predicted subjective re-experiencing ratings, we fitted a linear mixed-effect model using the lme4 ^82^, with re-experiencing ratings as the dependent variable, EMG amplitude, encoding condition and their interaction as fixed effects, and random intercepts for participants and events. To assess whether performance in identifying the encoding condition differed from chance level, we performed a one-sample t-test against chance (0.5). All tests were two-tailed with α = 0.05.

