## Supplementary Material for "Re-membering: spontaneous reactivation of motor cortex during memory re-experiencing"

1.1. Supplementary material
     1.1.1. Experimental Paradigm supporting information

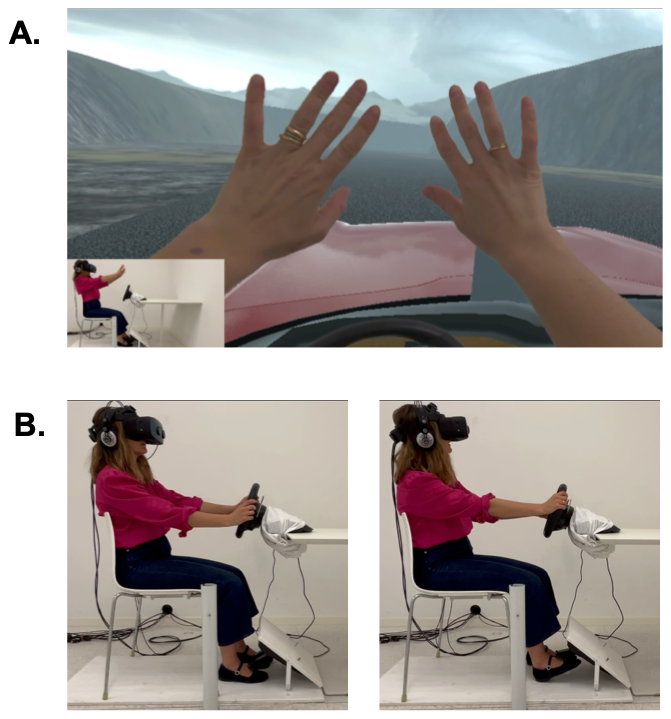
**Supplementary Figure 1: Encoding mixed-reality set-up. A,** Participants were immersed in a virtual environment where they could see their real body in real time. **B,** Participants were equipped with a mixed-reality headset and seated in a white room in front of a steering wheel and with a foot pedal above their right foot. During “foot” events, participant lifted the foot pedal with their right foot (right panel), which otherwise was at rest (left panel). Photographs are of the authors.

**Free retrieval instructions**

*“A word cue will appear on the screen, indicating the name of the obstacle to remember.*

*When the word disappears, please close your eyes, and try to remember as much as you can from the moment you saw this obstacle.*

*Try to remember the obstacle moving in front of you and all the details of the scene, but also try to imagine yourself in the scene again, as if you were travelling back in time to re-experience or relive the scene.”*

**Supplementary Box 1:** Standardized free re-retrieval instructions

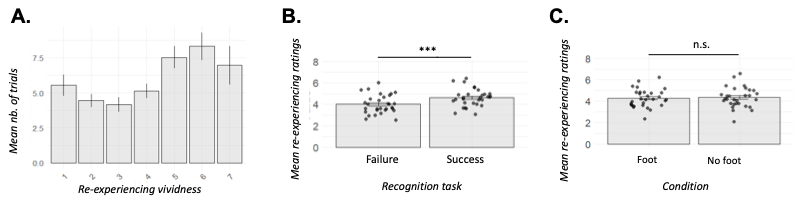
**Supplementary Figure 2:** Re-experiencing in Study 1. **A,** Distribution of re-experiencing ratings across the 7 levels of the scale (1-Not re-experiencing; 7-Fully re-experiencing). Error bars correspond to standard error of the mean across subjects. **B,** Re-experiencing ratings were higher when participants successfully recognized the obstacle in the recognition task. Each dot represents a single participant. **C,** Re-experiencing ratings did not differ between foot and no foot events. *** p<0.001, n.s.: non significant.

**Obstacle names**

*(English translation)*

Accident

Haystack

Bushes

Bus

Shopping carts

Boxes

Dog

Cones

Drone

Moose

Kite

Foodtruck

Smoke

Crane

Balloon man

Gardener

Rabbit

Laundry

Stilt house

Windsock

Hot air balloon

Sheep

Birds

Ad board

Police

Drawbridge

Gate

Paint

Waste bins

Rocks

Road roller

Pile of leaves

Lawn mower

Barrels

Train

Skate boarder

Scooter

Leaky pipe

Cyclist

Old man

**Supplementary Table 1:** Full obstacle list with obstacle names in English for the purpose of this article (in the original version of the task obstacle names were in French).

1.1.2. Connectivity analysis – Generalized psychophysiological interaction

| **Seed** | **Cluster** | | **Peak MNI coordinates** | | |
| --- | --- | --- | --- | --- | --- |
|  | **Size** | **pFWE** | **x** | **y** | **z** |
| Left HPC | 3624 | 0,00 | -32 | -48 | -42 |
|  | 1545 | 0,00 | 12 | -68 | 52 |
|  | 237 | 0,00 | -4 | 14 | 50 |
|  | 188 | 0,00 | 28 | 2 | 70 |
|  | 168 | 0,00 | -16 | -26 | 1 |
|  | 149 | 0,00 | 22 | -58 | -8 |
|  | 130 | 0,00 | 46 | 32 | 30 |
|  | 118 | 0,00 | -24 | -48 | -14 |
|  | 105 | 0,00 | -48 | -36 | 0 |
|  | 89 | 0,00 | 2 | -66 | 8 |
|  | 63 | 0,00 | 12 | 10 | 14 |
|  | 51 | 0,01 | -18 | -4 | -22 |
|  | 49 | 0,02 | 60 | -56 | 22 |
| Right HPC | 1830 | 0,00 | -12 | -94 | -8 |
|  | 1036 | 0,00 | -48 | -32 | 54 |
|  | 859 | 0,00 | 4 | -56 | -20 |
|  | 312 | 0,00 | 10 | -68 | 50 |
|  | 179 | 0,00 | -26 | -50 | 42 |
|  | 130 | 0,00 | 44 | -36 | 42 |
|  | 72 | 0,00 | 18 | -70 | -24 |
|  | 70 | 0,00 | 30 | -70 | 28 |
|  | 69 | 0,00 | -40 | -54 | -36 |
|  | 64 | 0,00 | 12 | -62 | 50 |
|  | 52 | 0,01 | -26 | -36 | -42 |

***Supplementary Table 2:*** *gPPI seed-to-whole brain analysis on free retrieval results with left (upper table) and right (lower table) hippocampal seeds.*

1.1.3. Imaging results – Re-experiencing Parametric Modulation

| **Label** | **Peak MNI coordinates** | | | **Cluster size** | **T value** |
| --- | --- | --- | --- | --- | --- |
|  | **x** | **y** | **z** |  |  |
| right inferior frontal gyrus | 42 | 4 | 40 | 152 | 5,18 |
| right frontal middle gyrus | 50 | 36 | 8 | 107 | 4,81 |
| left SMA / right SMA | 4 | 0 | 68 | 138 | 4,60 |
| right temporal middle gyrus | 54 | -58 | 14 | 89 | 4,45 |
| right angular gyrus / right occipital middle gyrus | 42 | -72 | 28 | 140 | 4,32 |
| left supramarginal gyrus | -60 | -36 | 28 | 117 | 4,15 |

**Supplementary Table 3:** Clusters showing significant positive modulation of BOLD activity by re-experiencing ratings.
